# Species-specific effects of cation channel TRPM4 small-molecule inhibitors

**DOI:** 10.1101/2021.06.01.446517

**Authors:** Prakash Arullampalam, Barbara Preti, Daniela Ross-Kaschitza, Martin Lochner, Jean-Sébastien Rougier, Hugues Abriel

**Affiliations:** Institute of Biochemistry and Molecular Medicine, and Swiss National Centre of Competence in Research (NCCR) TransCure, University of Bern, Bühlstrasse 28, 3012, Bern, Switzerland

**Keywords:** TRP channels, TRPM4, 9-phenanthrol, comparative physiology, mouse models, patch-clamp

## Abstract

**Background:** The Transient Receptor Potential Melastatin member 4 (*TRPM4*) gene encodes a calcium-activated non-selective cation channel expressed in several tissues. Mutations in *TRPM4* have been reported in patients with different types of cardiac conduction defects. It is also linked to immune response and cancers, but the associated molecular mechanisms are still unclear. Thus far, 9-phenanthrol is the most common pharmacological compound used to investigate TRPM4 function. We recently identified two promising aryloxyacyl-anthranilic acid compounds (abbreviated CBA and NBA) inhibiting TRPM4. However, all afore-mentioned compounds were screened using assays expressing human TRPM4, whereas the efficacy on mouse TRPM4 has not been assessed. Mouse models are essential to investigate ion channel physiology and chemical compound efficacy.

**Aim:** In this study, we performed comparative electrophysiology experiments to assess the effect of these TRPM4 inhibitors on human and mouse TRPM4 channels heterologously expressed in TsA-201 cells.

**Methods and Results:** We identified striking species-dependent differences in TRPM4 responses. NBA inhibited both human and mouse TRPM4 currents when applied intracellularly and extracellularly using excised membrane patches. CBA inhibited human TRPM4, both intracellularly and extracellularly. Unexpectedly, the application of CBA had no inhibiting effect on mouse TRPM4 current when perfused on the extracellular side. Instead, it increased mouse TRPM4 current at negative holding potentials. In addition, CBA on the intracellular side altered the outward rectification component of the mouse TRPM4 current. Application of 9-phenanthrol, both intracellularly and extracellularly, inhibited human TRPM4. For mouse TRPM4, 9-phenanthrol perfusion led to opposite effects depending on the site of application. With intracellular 9-phenanthrol, we observed a tendency towards potentiation of mouse TRPM4 outward current at positive holding potentials.

**Conclusion:** Altogether, these results suggest that pharmacological compounds screened using “humanised assays” should be extensively characterised before application in *in vivo* mouse models.

## Introduction

Transient Receptor Potential (TRP) ion channels constitute a superfamily of cationic channels permeable to Na+, Ca^2+^, and/or Mg2+ (Nikolaev et al., 2019). By contributing to intracellular Ca^2+^ signalling (Clapham et al., 2001;Montell et al., 2002), they are critically involved in various calcium-dependent cell functions. The TRP family member TRPM4 has received much attention due to its implication in many physiological functions such as cardiac activity (Guinamard and Bois, 2007;Liu et al., 2010;Stallmeyer et al., 2012;Guinamard et al., 2015;Piao et al., 2015), immune response (Launay et al., 2004;Barbet et al., 2008;Hemmer et al., 2015), cancer (Sagredo et al., 2018), arterial constriction (Earley et al., 2004), insulin secretion (Cheng et al., 2007), and cell death (Simard et al., 2012). TRPM4 is expressed in many organs, including the heart, kidney, and brain (Reading and Brayden, 2007;Abriel et al., 2012;Simard et al., 2013;Flannery et al., 2015;Mehta et al., 2015), and is upregulated in specific cancer cells, rendering it a promising therapeutic target (Berg et al., 2016;Gao and Liao, 2019;Sagredo et al., 2019). Unlike most other TRP channels, which are calcium-permeable, TRPM4 is only permeable to monovalent cations, while intracellular calcium activates the channel in a voltage-dependent manner. Its activation leads to plasma membrane depolarisation, thereby regulating calcium homeostasis by decreasing the driving force for calcium to enter the cell through calcium-permeable channels (Launay et al., 2002;Nilius et al., 2004).

Many pre-clinical *in vivo* pharmacological studies are performed in mouse models, but TRPM4-inhibiting compounds are generally screened using heterologous systems expressing the human TRPM4 channel. It is highly relevant to study the effect of TRPM4-inhibiting compounds on human and mouse TRPM4, as TRP channel orthologues from different species may show remarkable functional differences: Mammalian TRPV1, for instance, is activated by capsaicin, whereas chicken trpv1 is entirely insensitive for it (Jordt and Julius, 2002;Chu et al., 2020). Another example is the species-dependent effect of CMP1, a thioaminal-containing compound that activates rat TRPA1 but blocks human TRPA1 (Chen et al., 2008). Similarly, caffeine activates mouse TRPA1, but suppresses human TRPA1 (Nagatomo and Kubo, 2008). Other studies have shown species-specific effects of protein modulators and temperature on TRPM8 (Hilton et al., 2018) and TRPV1 (Papakosta et al., 2011), respectively.

In this study, we characterised the specificity and potency of three TRPM4 inhibitors (Fig. 1A): 9-phenanthrol, 4-chloro-2-[2-(2-chloro-phenoxy)-acetylamino]-benzoic acid (CBA, formerly named compound 5 in (Ozhathil et al., 2018)), and 4-chloro-2-(2-(naphthalene-1-yloxy) acetamido) benzoic acid (NBA, formerly named compound 6 in (Ozhathil et al., 2018)). Compound 9-phenanthrol has been widely used to inhibit TRPM4 in functional assays. However, its poor selectivity limits the validity of its use in animal models or primary cell lines (Burris et al., 2015;Ma et al., 2017). It is particularly problematic that past studies on TRPM4 have based their conclusions on patch-clamp assays that used this inhibitor despite its low potency and low specificity. CBA and NBA have been developed by our group and more potently inhibit TRPM4 than 9-phenantrol does (Ozhathil et al., 2018). Since these pharmacological compounds were discovered in a screening assay based on cell lines expressing human TRPM4, however, it was crucial to study their effect on mouse TRPM4.

**Figure 1.**
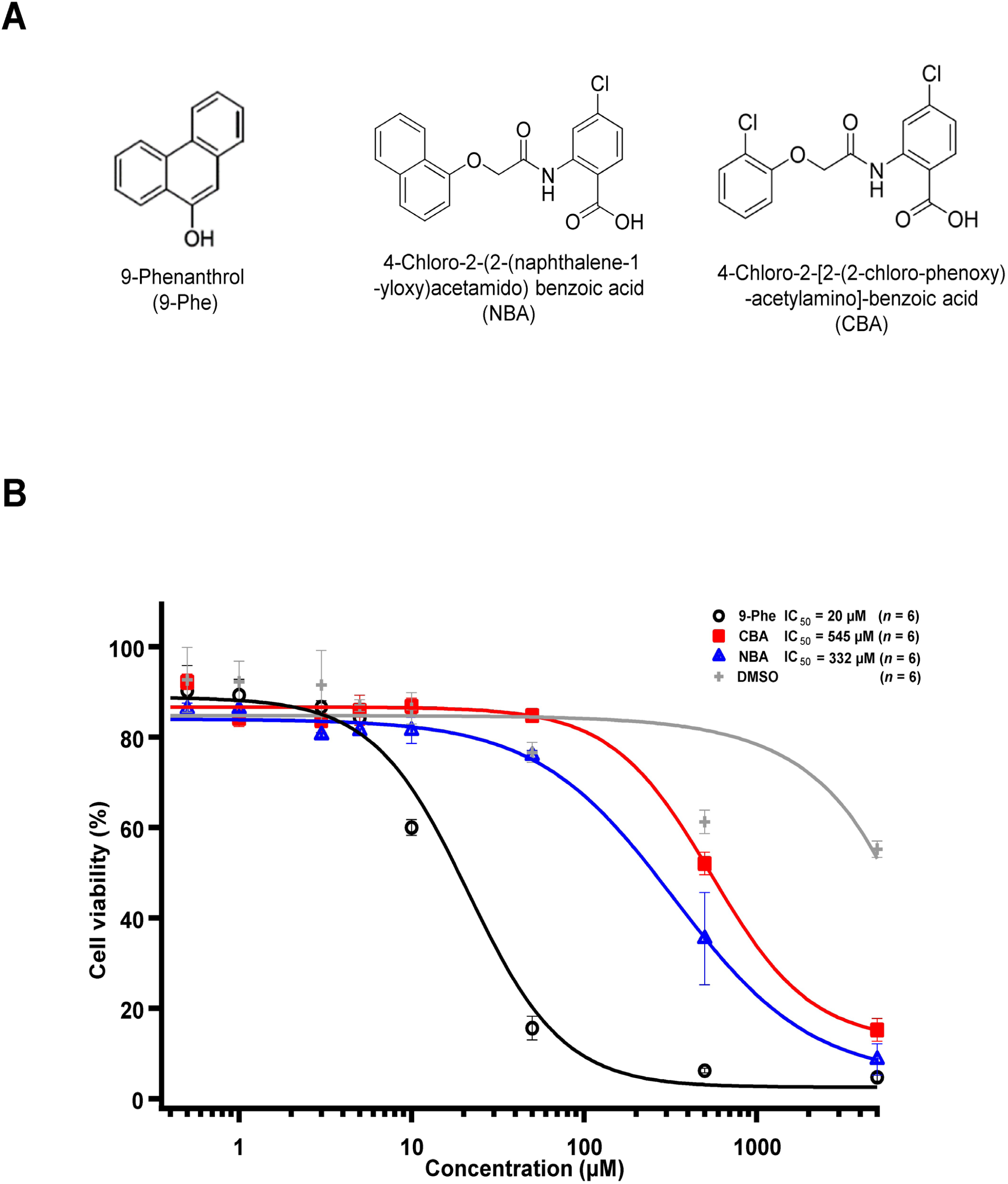
TRPM4 pharmacological compounds used in this study. (A) Structure of 9-phenanthrol (left), NBA (middle), and CBA (right). (B) Concentration-response curve showing the cytotoxicity of 9-phenanthrol (black circles), CBA (red squares), NBA (blue triangles), and DMSO (grey plus symbols) (*n* = 6 for each concentration).

Using the patch-clamp technique in cells expressing human or mouse TRPM4, we characterised the differences in specificity and potency of CBA, NBA, and 9-phenanthrol. We observed that CBA inhibits human TRPM4 but not mouse TRPM4, while NBA inhibits mouse and human TRPM4. Finally, 9-phenanthrol inhibits human TRPM4 and affects mouse TRPM4 currents differently depending on whether it is applied on the intracellular or extracellular side.

## Materials and Methods

### Cell culture

Transformed human embryonal kidney (TsA-201) cells stably overexpressing either mouse TRPM4, human TRPM4, or empty vector (pCDNA3.1 Zeo (+)) were generated using zeocin as a selection antibiotic. The cells were cultured in Dulbecco’s Modified Eagle’s medium (DMEM, #41965039) (Invitrogen, Zug, Switzerland) supplemented with 10% FBS, 4 mM glutamine, penicillin/streptomycin (50 U/ml), and Zeocin® 1/500 (Invitrogen cat# R250-01). They were kept at 37°C with 5% CO2 in a humidified atmosphere incubator.

### Western blot

To detect TRPM4 protein expression, whole-cell lysates were prepared by lysing cells in lysis buffer [50 mM HEPES pH 7.4; 150 mM NaCl; 1.5 mM MgCl_2_, 1 mM EGTA pH 8.0; 10% glycerol; 1% Triton X-100; complete protease inhibitor cocktail (Roche, Mannheim, Germany)] for 1 h at 4°C. Cell lysates were centrifuged for 15 min at 16 000 × g at 4°C, pellets discarded, and protein concentration of the supernatant was evaluated using the Bradford assay. Sixty μg of each protein sample was run on 9% polyacrylamide gels and transferred to a nitrocellulose membrane with the Turbo Blotdry blot system (Biorad, Hercules, CA, USA). After transfer, membranes were blocked with 0.1% BSA in PBS and incubated with rabbit primary anti-TRPM4 antibodies (epitope: VGPEKEQSWIPKIFRKKVC) (generated by Pineda, Berlin, Germany) diluted 1:750 in PBS/Tween using the SNAP i.d. system (Millipore, Billerica, MA, USA), followed by incubation with secondary antibodies (IRDye 800CW 1:1000 in PBS/Tween, LI-COR Biosciences, Lincoln, NE, USA). Membranes were scanned using the LiCor Odyssey Infrared imaging system (LI-COR Biosciences).

### Compounds tested

9-phenanthrol (9-Phe) was obtained from Sigma-Aldrich (Buchs SG, Switzerland). Compounds CBA, NBA, and the other anthranilic acid amides (Fig. 7) were synthesised as previously described (Ozhathil et al., 2018).

### Inside-out patch clamp

Electrophysiological recordings were performed in the inside-out patch-clamp configuration with patch pipettes (1–2 μm tip opening) pulled from 1.5 mm borosilicate glass capillaries (Zeitz-Instruments GmbH, München, Germany) using micropipette puller P 97 (Sutter Instruments, Novato, CA, USA). The tips were polished to have a pipette resistance of 2–4 MΩ in the bath solution. The pipette solution contained (in mM) 150 NaCl, 10 HEPES, and 2 CaCl_2_ (pH 7.4with NaOH). The bath solution contained (in mM) 150 NaCl, 10 HEPES, 2 HEDTA (pH 7.4 with NaOH), and no Ca^2+^. Solutions containing 300 μM Ca^2+^ were made by adding the appropriate concentration of CaCl_2_ without buffer to a solution containing (in mM) 150 NaCl, 10 HEPES (pH 7.4 with NaOH) as reported previously (Zhang et al., 2005). Bath solution with 0 and 300 μM Ca^2+^ concentrations were applied to cells by a modified rapid solution exchanger (Perfusion Fast-Step SF-77B; Warner Instruments Corp., CT, USA). Membrane currents were recorded with a Multiclamp 700B amplifier (Molecular Devices, Sunnyvale CA, USA) controlled by Clampex 10 via Digidata 1332A (Molecular Devices, Sunnyvale, CA, USA). Data were low-pass filtered at 5 kHz and sampled at 10 kHz. Experiments were performed at room temperature (20–25°C). The holding potential was 0 mV. For measuring steady-state currents, the stimulation protocol consisted of two sweeps with a total duration of 1000 ms, the first sweep at -100 mV for 500 ms and the second at +100 mV for 500 ms (Fig. 2). The effect of the compounds on TRPM4 current has been calculated by averaging the last 100 ms of the second sweep at +100 mV. The I-V relationships (IV curves) have been determined using an I-V protocol from -100 mV to +100 mV for 500 ms with a difference of 20 mV between each sweep (cf. Fig. 6). The IV curves have been normalised to the maximum current. No fit was applied.

**Figure 2.**
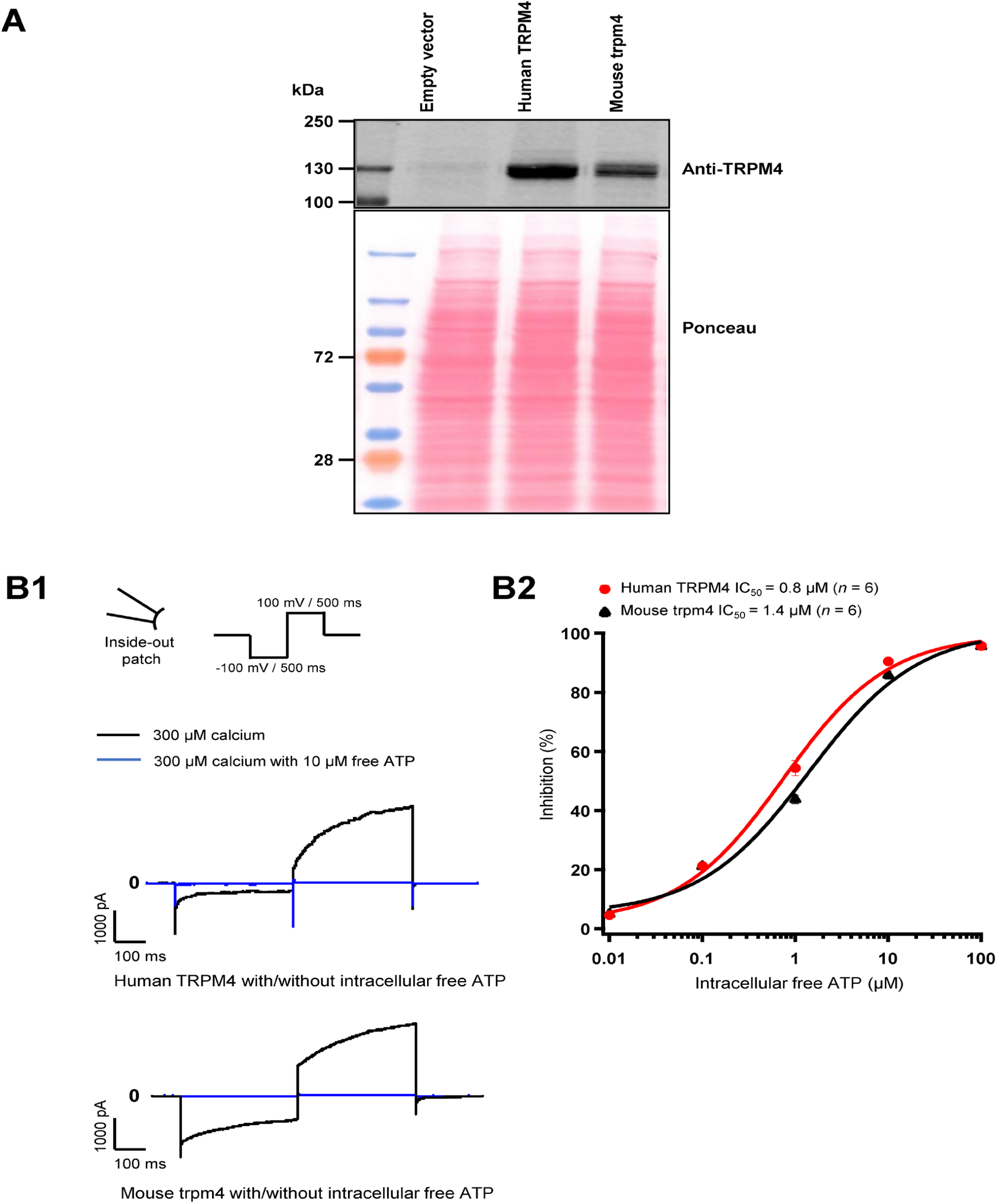
Validation of cell lines stably expressing TRPM4. (A) Western blot showing TRPM4 expression as detected by anti-TRPM4 antibody from whole-cell extracts prepared from TsA-201 cells. (B) Effect of free ATP on TRPM4 current. B1, representative traces of TRPM4 current activated by 300 μM calcium (black) and inhibited by 10 μM free ATP (blue) currents recorded from excised membrane patches (inside-out) (up and lower panels, respectively, human and mouse TRPM4). B2, the effect of free ATP on TRPM4 current. The values were fitted with the Hill equation to interpolate IC50 values of free ATP (*n* = 6 for each concentration).

### Application of the compounds and ATP

After obtaining inside-out membrane patches, the presence of TRPM4 in the patch membrane was confirmed by switching the perfusion solution without calcium at the intracellular side of the membrane to a solution containing 300 μM calcium, which activates TRPM4 and triggers the current. Once the TRPM4 currents were activated and stabilised for intracellular compound application, the perfusion solution containing 300 μM calcium and either the compound or free adenosine triphosphate (ATP) was applied. For extra-cellular application, the compounds were added to the intracellular (pipette) solution.

### Cytotoxicity test

TsA-201 cells were seeded in a 96-wells plate at a concentration of 10 000 cells per well with DMEM and incubated at 37°C and 5% CO2 in a humidified atmosphere incubator overnight. After reaching 80% to 90% of cell confluence, the cells in the 96-well plate were treated with different compound concentrations (0.5, 1, 2, 4, 10, 50, 500, and 5000 µM of CBA, NBA, and 9-phenanthrol) for 24 hours. After incubation, the cells were washed and incubated in 0.01% trypsin-EDTA until the cellular layer was completely detached. Cells were centrifuged down, and the pellet resuspended in 1 mL PBS. Cells were then diluted in 1 part of 0.4% trypan blue and 1 part cell suspension. This mixture was incubated for 3 min at room temperature. Finally, the unstained (viable) and stained (nonviable) cells were counted in the haemocytometer (BRAND® counting chamber BLAUBRAND®, Wertheim, Germany).

### Data analyses and statistics

Electrophysiology data were analysed using IGOR PRO 6 (WaveMetrics, Lake Oswego, USA), Normalized current percentage was calculated as follows: ([activated current at 300 μM calcium] – [activated current at 300 μM calcium with compound]) * 100 / [activated current at 300 µM calcium]. Concentration-response curves were fitted using the Hill equation fit parameter of IGOR (NC = NC_max_ [Cmpd]nH/([Cmpd]nH + EC_50_ nH)) where NC is normalized current, [Cmpd] is compound concentration and nH is the Hill coefficient. Data are presented as mean ± SEM except for Figures 3 and 7, where the data have been normalised to 100 for the current elucidated in the presence of 300 µM calcium only. Statistically significant differences conditions were determined using a non-parametric test with a Mann-Whitney post-test. *P*-values ≤ 0.01 were considered as statistically significant and marked with ** in respective figure panels.

## Results

### Nine-phenanthrol is more cytotoxic than CBA and NBA

First, we tested the cytotoxicity of the three selected TRPM4 compounds (Fig. 1A). The calculated EC_50_, corresponding to the concentration of the inhibitor required to induce 50% of cell death, was 15- to 25-fold lower for 9-phenanthrol (EC_50_ ∼ 20 μM) than for NBA (EC_50_∼ 332 μM) and CBA (EC_50_ ∼ 545 μM), respectively (Fig. 1B). These observations illustrate that the two new compounds CBA and NBA, are less cytotoxic than 9-phenanthrol.

### Characterisation of TsA-201 cell lines stably overexpressing either human or mouse TRPM4

To characterise the TsA-201 cell lines stably overexpressing either wild-type human or mouse TRPM4 generated for this study, we first tested the TRPM4 protein expression by western blot (Fig. 2A). Using an antibody recognising TRPM4 from both species, a robust expression of TRPM4 was observed in these modified cells compared to the cells transfected with an empty vector (Fig. 2A). To investigate if these cell lines express functional channels, we performed inside-out patch-clamp experiments to first record the current and second to investigate the well-known TRPM4 inhibitory effect of free ATP on TRPM4 currents (Fig. 2B1 and 2B2). As expected, robust TRPM4 currents were observed in TsA-201 cell lines stably expressing either human or mouse TRPM4 channels compared to untransfected cells (data not shown). This current decreased after intracellular application of free ATP (Fig. 2B1 and 2B2). On the concentration-response curves depicted in Fig. 2B2, the intracellular ATP IC_50_ for both types of channels was 0.8 μM for human TRPM4 and 1.4 μM for mouse TRPM4 (Fig. 2B2). Altogether, these results confirm that both cell lines expressed functional human or mouse TRPM4 wild-type channels and can be used for further investigations.

### Nine-phenanthrol inhibits human TRPM4 current and upregulates mouse TRPM4 current when applied intracellularly

Knowing that lipophilicity tools have predicted that 9-phenanthrol may cross the plasma membrane, we decided to record mouse and human TRPM4 currents in the presence of the three compounds (9-phenanthrol, CBA, and NBA) either applied at the intracellular side (bath solution) or the extracellular side (pipette solution).

First, we tested the effects of 9-phenanthrol on human and mouse TRPM4 channels. Thirty μM 9-phenanthrol, corresponding to the previously determined IC_50_ (Fig. 1B), was applied using a perfusion solution containing 300 μM calcium (Ozhathil et al., 2018). Nine-phenanthrol inhibited human TRPM4 sodium current with a similar efficacy independently of the application side (extracellular or intracellular) (Fig. 3A1, 3B1, 3A2, and 3B2). Unexpectedly, while extracellular (pipette solution) 9-phenanthrol decreased the mouse TRPM4 current, it increased the current when applied intracellularly (Fig. 3A1, 3B1, 3A2, and 3B2). This increase was observed only in the presence of 300 μM calcium, suggesting that 9-phenanthrol acts as a potentiator rather than as an activator of the mouse TRPM4 current (Supplementary Fig. 1A). Moreover, this effect was more pronounced with low concentrations (0.1 μM) of 9-phenanthrol than with higher concentrations (Supplementary Fig. 1B). When applying 100 μM 9-phenanthrol, the observed potentiator effect was smaller than in the presence of 0.1 µM 9-phenanthrol (Supplementary Fig. 1B). Altogether, these data illustrate that the effects mediated by 9-phenanthrol depend on the application side of the excised membrane patch and are species-specific.

**Figure 3.**
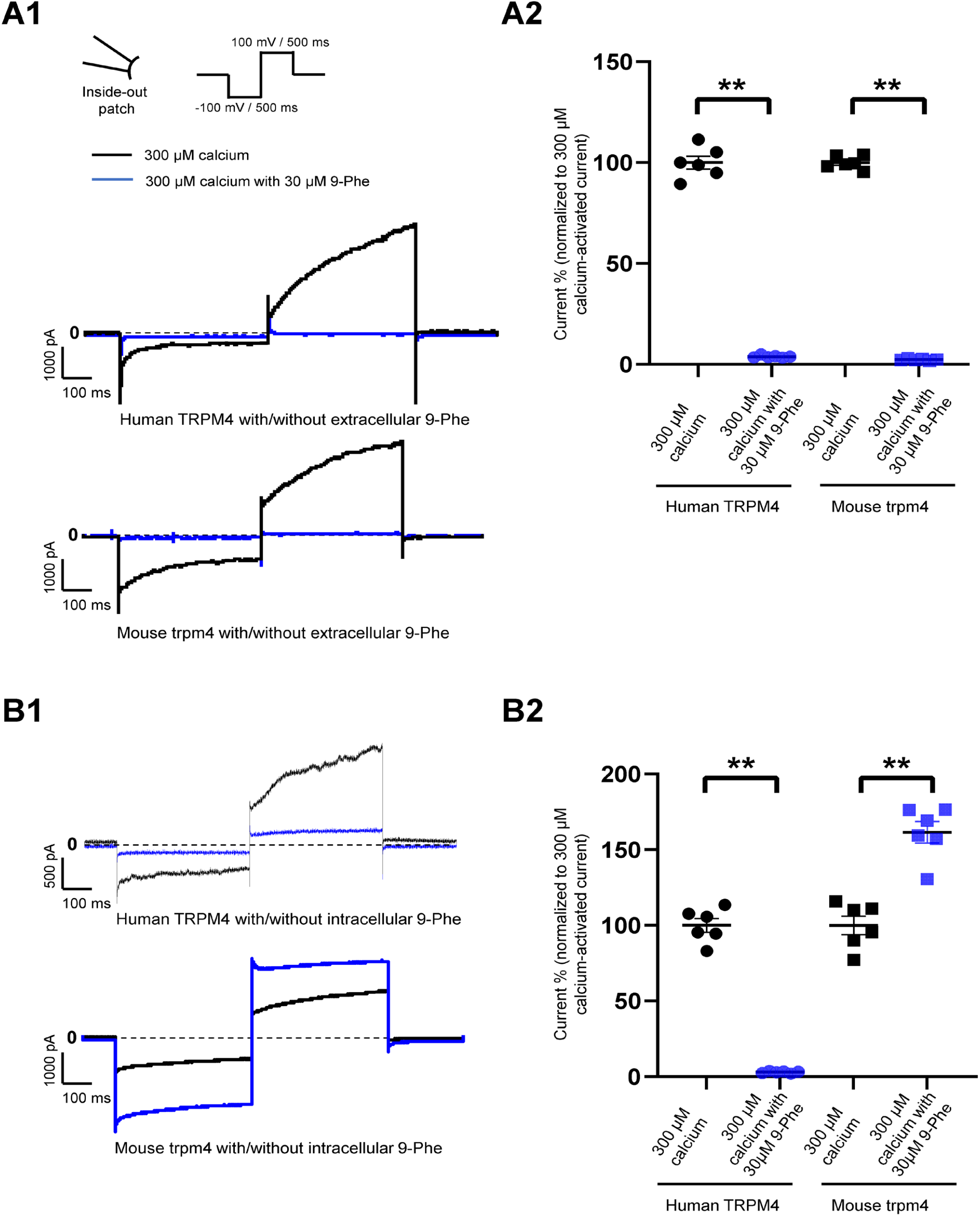
Species-specific effects of 9-phenanthrol on TRPM4 currents. Representative traces of TRPM4 currents recorded from excised membrane patches (inside-out) from cells expressing human (A1 and B1 upper panels) and mouse (A1 and B1 lower panel) TRPM4. Thirty-μM 9-phenanthrol is added to the intracellular (A) or extracellular (B) side (black traces: control and blue traces; treated). All TRPM4 currents were obtained after perfusion with 300 μM free calcium solutions. (A2 and B2) Human TRPM4 (circle) and mouse TRPM4 (square) average current amplitudes at the end of voltage step to +100 mV with (blue) or without (black) 9-phenanthrol. Voltage protocol is given at the top of panel A (*n* = 6). \*\**P* ≤ 0.01.

### NBA decreases both human TRPM4 and mouse TRPM4 currents

Analogous to our experiments with 9-phenanthrol, we performed patch-clamp recordings using the compound NBA. We observed that 10 μM NBA inhibited both human and mouse TRPM4 current independent of the side of the application (Fig. 4A1, 4A2, 4B1, and 4B2). The concentration-response curve for NBA revealed a sigmoid curve for both human and mouse TRPM4 currents (Fig. 4A2 and 4B2). Fitting the data with the Hill equation revealed that the IC_50_s of NBA are 0.187 μM and 0.119 μM for human and mouse TRPM4, respectively, when applied intracellularly (Fig. 4B2). The IC_50_s of extracellular NBA were 0.125 μM and 0.215 μM with human and mouse TRPM4, respectively (Fig. 4A2). Taken together, these data suggest that NBA does not discriminate between human and mouse TRPM4 channels and inhibits the TRPM4 current when applied to either side of the membrane.

**Figure 4.**
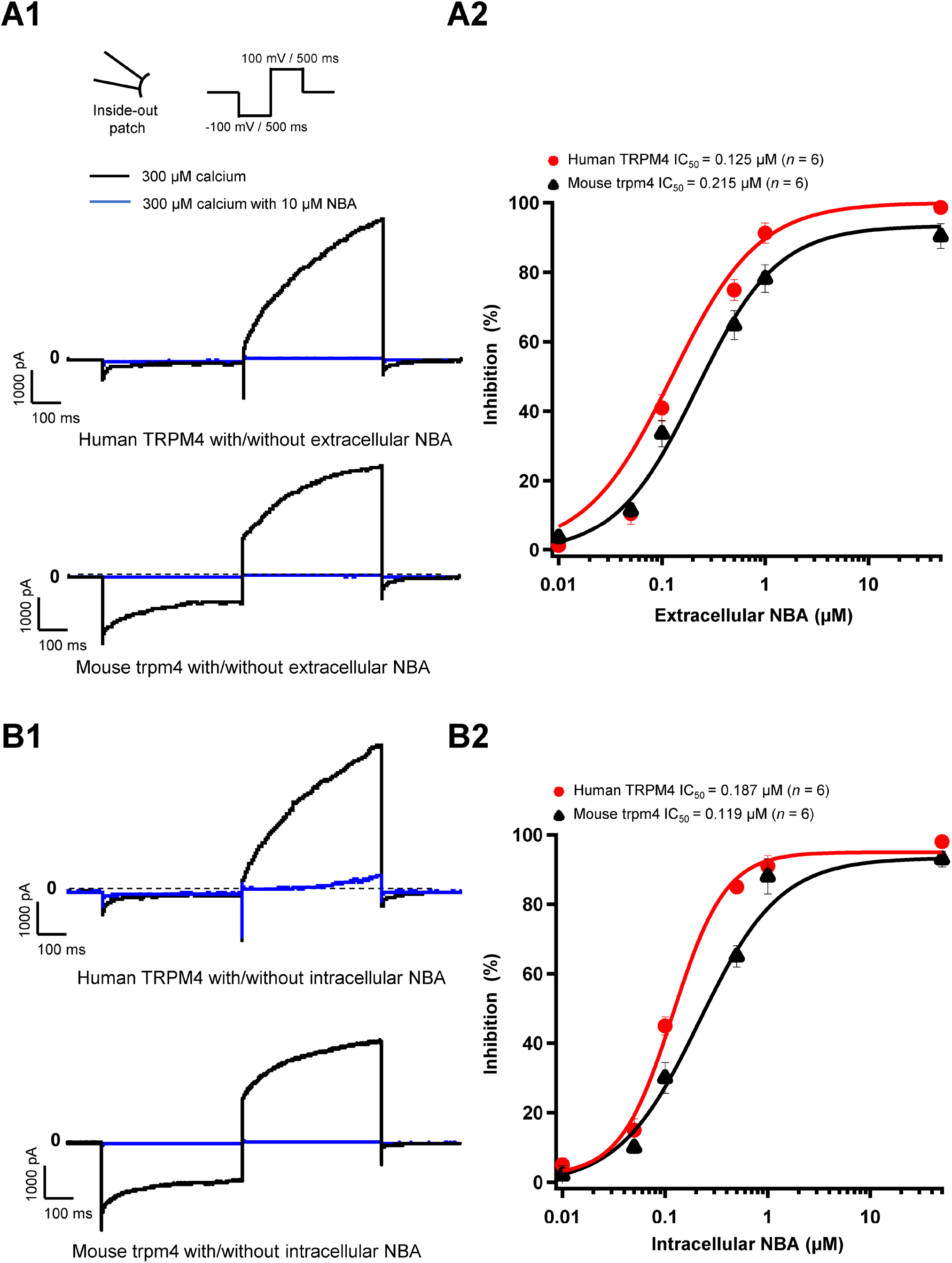
NBA has similar effects on human and mouse TRPM4 currents. (A1 and B1) Representative traces of TRPM4 currents recorded from excised membrane patches (inside-out). Human TRPM4 (A1, upper panel) and mouse TRPM4 (A1, lower panel) currents in the presence (blue) or absence of 10 μM NBA (black) in the pipette solution (extracellular application). Human (B1, upper panel) and mouse (B1, lower panel) TRPM4 currents in the presence (blue) or absence (black) of 10 μM NBA in the bath solution (intracellular application). All TRPM4 currents were obtained after perfusion with 300 μM free calcium solution. (A2 and B2) Concentration-response curves of human (red circles) and mouse (black triangles) TRPM4 for NBA either in pipette solution (extracellular application) (B1) or in the bath solution (intracellular application) (B2) (*n* = 6 for each concentration). Data were fitted with a Hill equation to interpolate the IC_50_.

### CBA decreases only human, not mouse, TRPM4 currents

Next, we assessed the effects of CBA on human and mouse TRPM4 currents. Application of 10 μM CBA decreased the human TRPM4 currents independent of the side of the application (Fig. 5A1, 5A2, 5B1, and 5B2). The IC_50_s were 0.7 μM (extracellular application/ pipette solution) and 0.8 μM (intracellular application) (Figs. 5A2 and 5B2). However, in membrane patches expressing mouse TRPM4 channels, neither the extracellular nor the intracellular application of CBA altered the TRPM4 currents (Fig. 5A1, 5A2, 5B1, and 5B2). Surprisingly, when CBA was applied intracellularly, a small but significant effect on mouse TRPM4 current was observed at positive voltages (+100 mV), while at negative membrane voltages (−100 mV), the current was significantly higher (Fig. 5B1 and Fig. 6). Based on this observation, we recorded a current-voltage relationship (I-V) protocol to quantify the effect of intracellular CBA application on mouse TRPM4 current at different membrane voltages (Fig. 6). As previously observed in figure 5, CBA increased the inward current at negative membrane voltages (Fig. 6) but only slightly altered the amplitude of the outward current at positives membrane voltages (Fig. 6). Doing so, CBA led to a linearisation of the I-V relationship removing the rectification features. Altogether, these results illustrate that the effect of CBA is species-specific, and CBA alters the inward current of mouse TRPM4.

**Figure 5.**
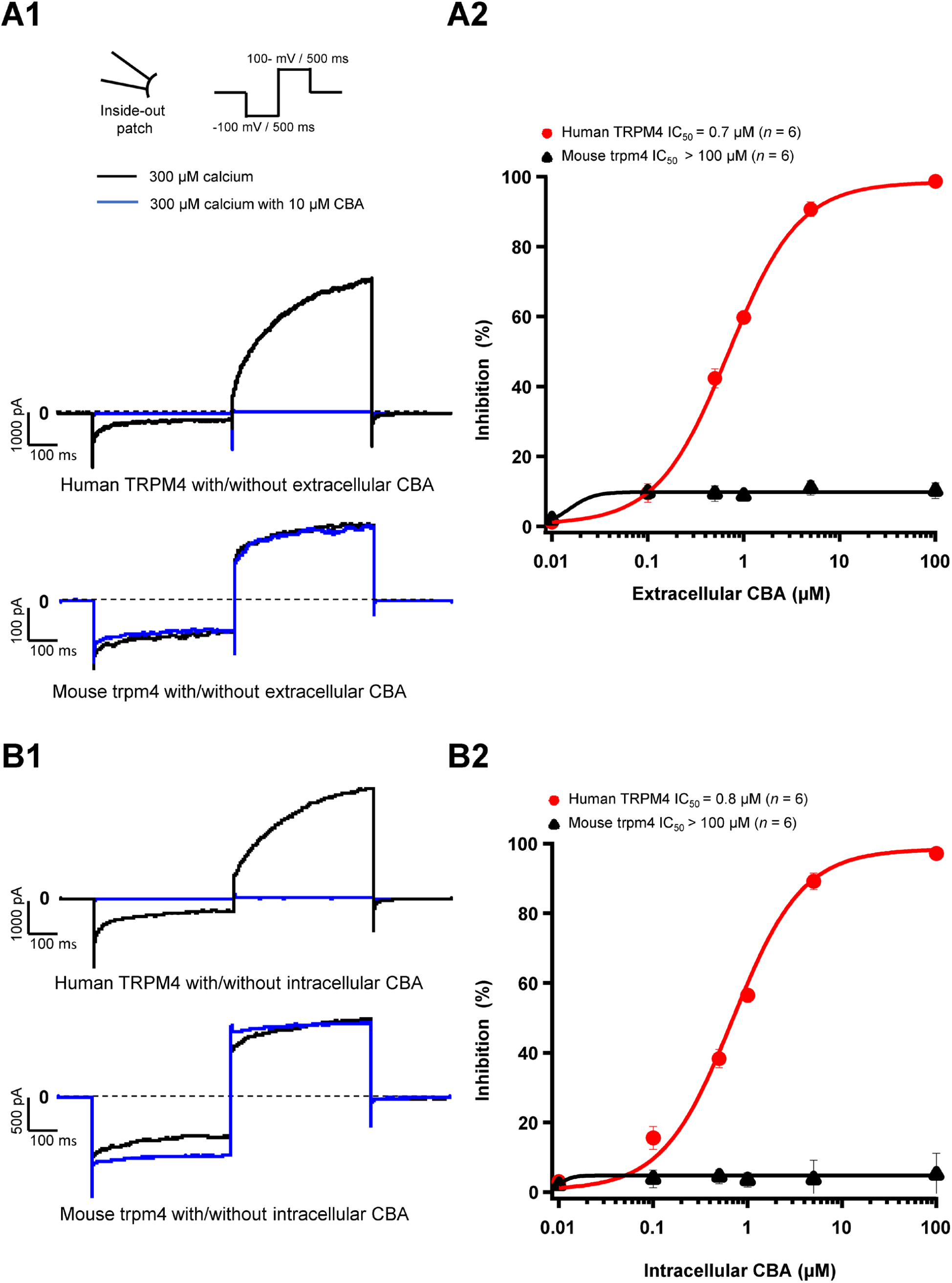
CBA blocks TRPM4 in a species-specific manner. (A1 and B1) Representative traces of TRPM4 currents recorded from excised membrane patches (inside-out). Human TRPM4 (A1, upper panel) and mouse TRPM4 (A2, lower panel) currents in the presence (blue) or absence (black) of 10 μM CBA in the pipette solution (extracellular application). Human (B1, upper panel) and mouse (B1, lower panel) TRPM4 currents in the presence (blue) or absence (black) of 10 μM CBA in the bath solution (intracellular application). All TRPM4 currents were obtained after perfusion with 300 μM of free calcium solution. (A2 and B2) Concentration-response curves on human (red circles) and mouse (black squares) TRPM4 currents for CBA. CBA was applied in the pipette solution (A2, extracellular application) or in the bath solution (B2, intracellular application) (*n* = 6 for each concentration). Data were fitted with a Hill equation to interpolate the IC_50_.

**Figure 6.**
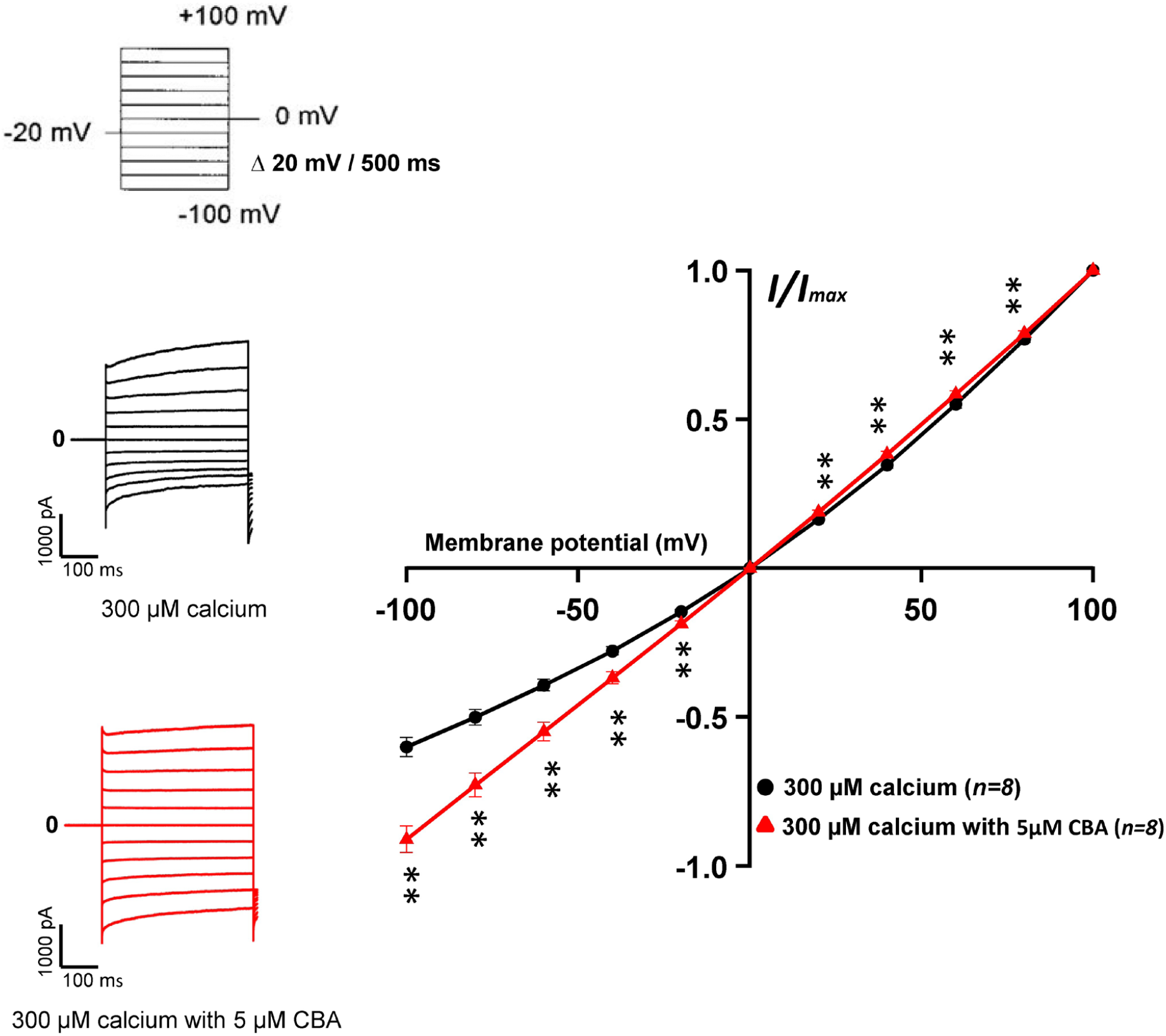
Effects of CBA on mouse TRPM4 current. Left panel shows raw traces of mouse TRPM4 currents recorded using the step protocol depicted at the top of the figure, with (black) or without (red) 5 μM CBA in the extracellular bath (intracellular application). Right panel shows IV curves, representing the average current amplitude at the end of each voltage step when the TRPM4 current was activated using 300 μM free calcium solution (black) and treated with 5 μM CBA (red) (*n* = 8). \*\**P* ≤ 0.01.

### NBA- and CBA-derived compounds have similar, but not the same, effects

To finally investigate if the effects observed could depend on further chemical structures, additional patch-clamp experiments were performed using compounds closely related to TRPM4 inhibitors CBA and NBA structures. They were selected based on a Structure-Activity Relationship (SAR) rationale (Ozhathil et al., 2018). The CBA-derived compound used was 4-chloro-2-(2-((4-chloro-2-methylphenoxy) propanamido) benzoic acid (CBA-d1) (Fig. 7A1). The selected NBA-derived compounds were 4-chloro-2-(2-((4-chloronaphthalen-1-yl) oxy) acetamido) benzoic acid (NBA-d1) and 4-chloro-2-(2-((5,6,7,8-tetrahydronaphthalen-1-yl) oxy) acetamido) benzoic acid (NBA-d2) (Fig. 7B1 and 7C1). Interestingly, similarly to CBA, CBA-d1 did not decrease the mouse TRPM4 currents when applied intracellularly (Fig. 7A1 and 7A2). Contrary to CBA-d1, NBA-d1 and NBA-d2 were able to inhibit mouse TRPM4 currents as also observed with NBA compounds (Fig. 7B1, 7B2, 7C1, and 7C2). Altogether, these results suggest that while some structurally related compounds behave similarly to the original one (NBA-d1 and NBA-d2 compared to NBA), some of them may have slightly different effects (CBA-d1 and CBA).

**Figure 7.**
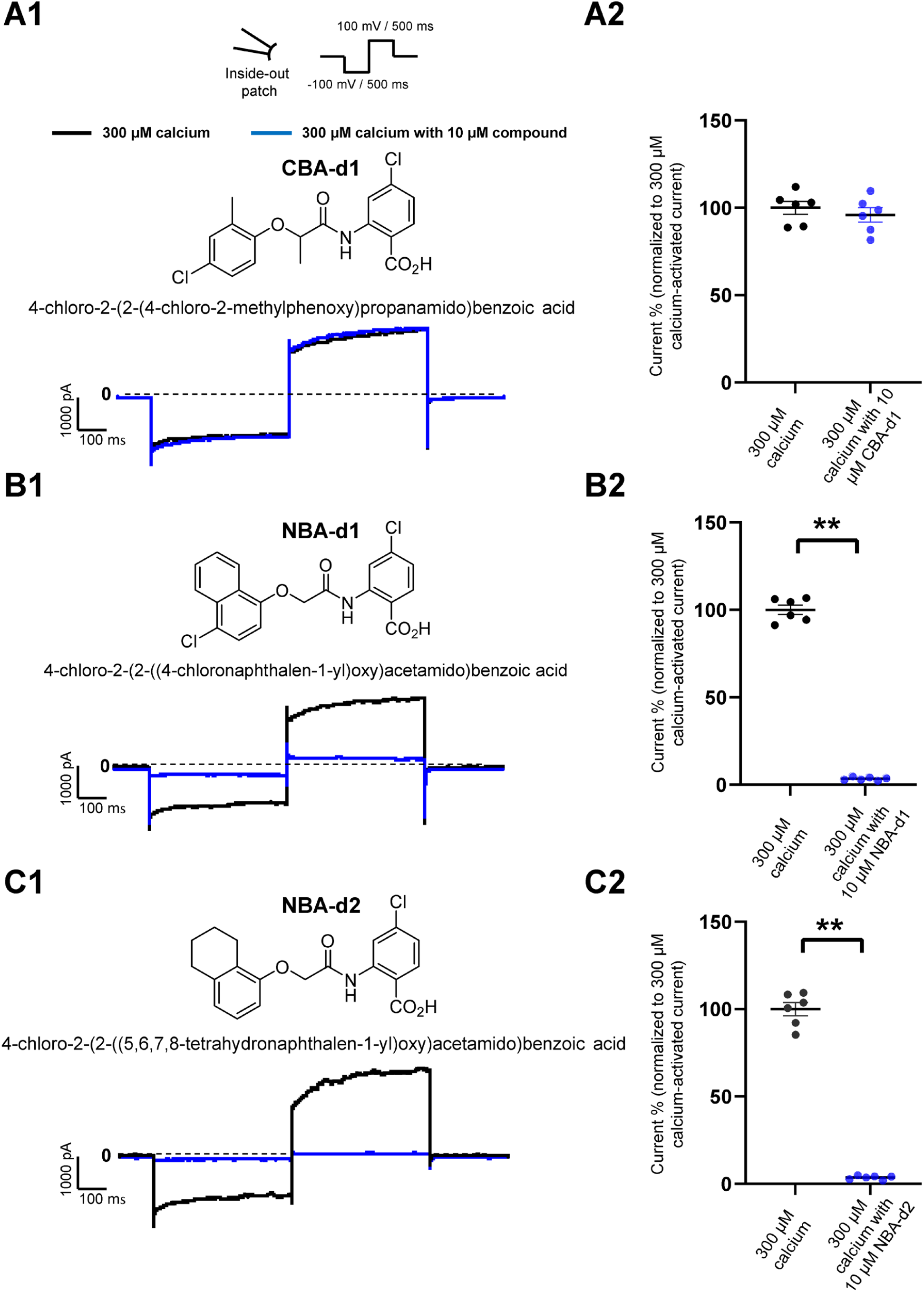
Effect of CBA- and NBA-derived compounds on mouse TRPM4 current. Chemical structures of pharmacological compounds derived from NBA and CBA (A1, B1, and C1; top panels). TRPM4 currents with 10 μM of either CBA-d1, NBA-d1, or NBA-d2 compounds. Mouse TRPM4 currents in the presence (blue) or absence (black) of 10 μM of CBA-d1 (A1 and A2), NBA-d1 (B1 and B2), and NBA-d2 (C1 and C2) in the bath solution (extracellular application). All currents were obtained after perfusion with 300 μM free calcium solution. (A2, B2, and C2) Average current amplitudes at the end of the voltage step to +100 mV. *n* ≥ 6 for all conditions. \**P* ≤ 0.05.

## Discussion

In this study, we assessed the species-specific effects of inhibitors of the calcium-activated non-selective monovalent ion channel TRPM4. Despite the relevance of TRPM4 for a wide range of physiological and pathological processes in different tissues, its roles remain not well understood. Identifying TRPM4 compounds that reliably block TRPM4 current in humans and mouse models is crucial to unravel the functions of TRPM4 and investigate it as a potential therapeutic target. We characterised the effect and species selectivity of three different TRPM4 small molecule inhibitors: 9-phenanthrol, NBA, and CBA. Further, we investigated three CBA and NBA derivatives. Nine-phenanthrol remains a widely used reference compound in studies to decipher the role of TRPM4 in mouse models. Although lipophilic prediction tools have suggested the capability of 9-phenanthrol to cross the plasma membrane, none of the studies published so far had investigated its effect when specifically applied on the intracellular side of the mouse TRPM4 channel using the inside out patch-clamp configuration. Here, we show for the first time that 9-phenanthrol inhibits mouse TRPM4 when applied on the extracellular part of the patch membrane, but its application on the inner membrane side leads to an evident potentiation of mouse TRPM4 current (Supplementary Fig. 1A). It remains unclear whether 9-phenanthrol has a direct effect on mouse TRPM4 or whether the observed increase in current is mediated by a second messenger such as Mg-ATP or phosphatidylinositol-4,5-bisphosphate, which are known to restore the desensitisation of TRPM4 current (Zhang et al., 2005). Further experiments are required to address this question. Considering that 9-phenanthrol may cross the plasma membrane altogether, these observations raised the question of the global effect on TRPM4 current of 9-phenanthrol would be when applied either in a mouse model or in cultured murine cells.

Interestingly, the potentiating effect of 9-phenanthrol when applied intracellularly to mouse TRPM4 was blunted at relatively high concentrations (Supplementary Fig. 1B). Although more experiments are required to decipher the molecular mechanisms behind these observations, it can be proposed that 9-phenanthrol, when applied at a high concentration at the intracellular part of the cell, can partly cross the lipid bilayer, reaching the extracellular part of the cell where it acts as an inhibitor. Conversely, 9-phenanthrol applied to the outside surface of cells may also cross the plasma membrane and cause a potentiating effect at the intracellular-facing surface. Based on these considerations, published studies that used 9-phenanthrol should be interpreted with caution.

Considering that (1) the IC_50_ to block TRPM4 is much higher for 9-phenanthrol than for NBA and CBA (Fig. 1), (2) 9-phenanthrol partially block other ion channels (Abriel et al., 2012), and (3) its cytotoxicity is higher than NBA and CBA, our results strongly suggest that 9-phenanthrol is a suboptimal tool to dissect the TRPM4 physiological role in mice. It is therefore recommended to perform future experiments on mouse TRPM4 with NBA instead of 9-phenanthrol. CBA has already been used in previous publications using cell lines expressing TRPM4 (Bianchi et al., 2018;Borgstrom et al., 2020). This compound is more potent and more specific than 9-phenanthrol concerning human TRPM4; indeed, CBA did not inhibit TRPV1, TRPV3, TRPV6, TRPM5, TRPM7, or TRPM8 (Ozhathil et al., 2018). CBA seems to be a promising inhibitor of human TRPM4, which is especially interesting for the cancer research field, where over-expression of human TRPM4 correlates with higher malignancy (Kappel et al., 2019;Borgstrom et al., 2020). CBA, however, does not inhibit the mouse TRPM4 current (Fig. 5). A possible explanation for this species-specific effect may lie in the 20% non-homology between mouse and human TRPM4 sequences. Generating chimeric constructs of mouse and human TRPM4 channels to address this hypothesis will be an important direction of study in the future.

Interestingly and unexpectedly, CBA alters biophysical features of the mouse TRPM4 (outward and inward currents) (Figs. 5B1 and 6), suggesting different mechanisms of action of CBA on the mouse and human TRPM4, the latter being entirely inhibited by CBA. The site(s) of CBA action on the mouse and human TRPM4 is/are not yet elucidated. The structure-function relationship explaining these observations thus remains unknown. CBA-derived compounds inhibit neither inward nor outward mouse TRPM4 currents, suggesting that the differences in their structure may affect the biophysical properties of the channel differently. Taken together, our results suggest that CBA should only be used to study functional properties of human TRPM4 and should be avoided in mouse models.

Finally, NBA and the structurally related compounds efficiently inhibit both mouse and human TRPM4 channels (Fig. 5). NBA is, therefore, a novel and promising tool compound to study the role of TRPM4 in both humans and mice. Nonetheless, its site(s) of action still need(s) to be explored. We propose that characterising the channel properties of TRPM4 in diverse animal species and identifying the molecular determinations for TRPM4 pharmacological compounds will provide a foundation to develop new drugs to target TRPM4. To better characterise these compounds, it will be interesting to perform similar experiments in human *in vitro* models using either primary (native) cells such as human cardiomyocytes or induced pluripotent stem cell-derived cardiomyocytes. Lastly, both CBA and NBA should be tested in TRPM4 orthologs from other species.

Previous characterisation by TRPM4-mediated Na+ influx assays and electrophysiology experiments has shown that NBA is about 4-9 times more potent than CBA (Ozhathil et al., 2018). Unfortunately, without a precise cryo-EM structure of the TRPM4 (Guo et al., 2017;Winkler et al., 2017) channel or other experimental evidence about the binding sites of CBA and NBA, it is challenging to explain why NBA is more potent than CBA. We can only speculate that the additional ring of the naphthalene substituent of NBA can form attractive interactions with a putative hydrophobic pocket of TRPM4, which the smaller 2-chlorophenyl ring of CBA cannot, or only to a much smaller degree. Molecular docking of CBA and NBA into human and mouse TRPM4 could help to suggest potential binding sites that need to be corroborated by follow-up mutagenesis studies.

Overall, the findings of the present study suggest that new compounds should be carefully characterised before used as tool compounds in animal models to avoid misinterpretation of the results. Based on our work, NBA is a promising tool compound to study the role of the TRPM4 channel in normal physiology and under pathological conditions. Unfortunately, the other compound, CBA, seems to be only interesting for experiments using human TRPM4 (Ozhathil et al., 2018). Finally, the TRPM4 inhibitor, 9-phenanthrol, which has been used for more than a decade to characterise TRPM4 function in different tissues, should be avoided as much as possible in favour of the two new characterised compounds NBA and CBA.

## Author Contributions

PA, DRK, and JSR conceived and designed the experiments. PA, BP, DRK, collected, analysed, and interpreted the data. PA, DRK, ML, JSR, and HA drafted the manuscript and revised it critically for important intellectual content.

## Acknowledgments

We thank Dr. Lijo Cherian Ozhathil for providing suggestions and feedback on the manuscript. We acknowledge Dr. Sarah Vermij for proofreading and editing.

## SUPPLEMENTARY MATERIAL

**Supplementary Figure 1.**
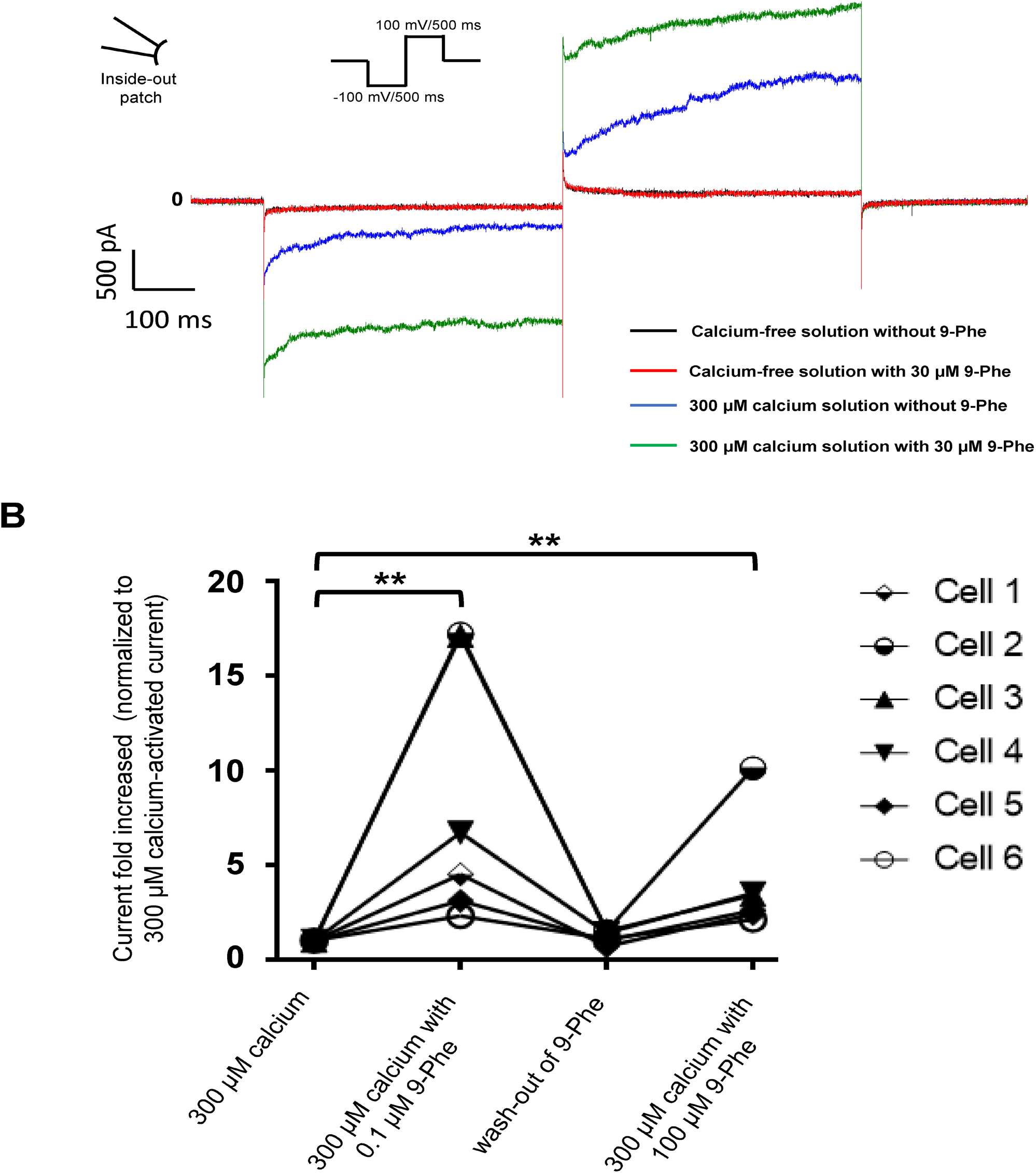
Nine-phenanthrol increases the mouse TRPM4 current only in the presence of calcium. **(A)** Representative traces of mouse TRPM4 currents recorded from excised membrane patches (inside-out) before and after application (intracellular application) of different solutions. Black trace: mouse TRPM4 current with a calcium-free and 9-phenanthrol-free solution; red trace: calcium-free and 9-phenanthrol solution with 30 μM of 9-phenanthrol; blue trace: 300 μM calcium solution without 9-phenanthrol; green trace: 300 μM calcium and 30 μM of 9-phenanthrol solution (similar observations were done with 3 different cells (data not shown). **(B)** Low concentrations of 9-phenanthrol potentiated more efficiently mouse TRPM4 currents compared to high concentrations. Normalised mouse TRPM4 currents from six different inside-out patches show a stronger potentiation of the current after low (0.1 μM) 9-phenanthrol concentration application (extracellular application) compared to a high (100 μM) concentration of this compound (*n* = 6).

